# Biophysical Basis of Paracellular Barrier Modulation by a Pan-Claudin-Binding Molecule

**DOI:** 10.1101/2024.11.10.622873

**Authors:** Chinemerem P. Ogbu, Alexandria M. Mandriota, Xiangdong Liu, Mason de las Alas, Srajan Kapoor, Jagrity Choudhury, Anthony A. Kossiakoff, Michael E. Duffey, Alex J. Vecchio

## Abstract

Claudins are a 27-member protein family that form and fortify specialized cell contacts in endothelium and epithelium called tight junctions. Tight junctions restrict paracellular transport across tissues by forming molecular barriers between cells. Claudin-binding molecules thus hold promise for modulating tight junction permeability to deliver drugs or as therapeutics to treat tight junction-linked disease. The development of claudin-binding molecules, however, is hindered by their intractability and small targetable surfaces. Here, we determine that a synthetic antibody fragment (sFab) we developed binds directly to 10 claudin subtypes with nanomolar affinity by targeting claudin’s paracellular-exposed surface. Application of this sFab to cells that model intestinal epithelium show that it opens the paracellular barrier comparable to a known, but application limited, tight junction modulator. This novel pan-claudin-binding molecule can probe claudin or tight junction structure and holds potential as a broad modulator of tight junction permeability for basic or translational applications.

## Introduction

Claudins are a family of ∼25 kDa membrane-embedded proteins that play the primary role in directing formation of tight junctions and regulating molecular transport through the paracellular spaces between individual cells within epithelial/endothelial sheets (**Fig. 1A**) *(1, 2)*. The human genome encodes for 27 individual claudin subtypes (*1*). Although claudins express in all mammalian tissues, individual subtypes have varied and tissue-specific expression patterns and levels that occur at different timepoints throughout tissue development (*3*). Additionally, at the molecular level, claudins display unique homo- and hetero-oligomeric compatibilities with other subtypes (*4*). This complex yet coordinated interaction network of claudins that constitute a given tissue thus directs its molecular transport properties—as claudin subtype incidence and variety influences the magnitude and morphology of tight junction strands—creating more or less leaky paracellular barriers (*1, 5*).

**Fig. 1.**
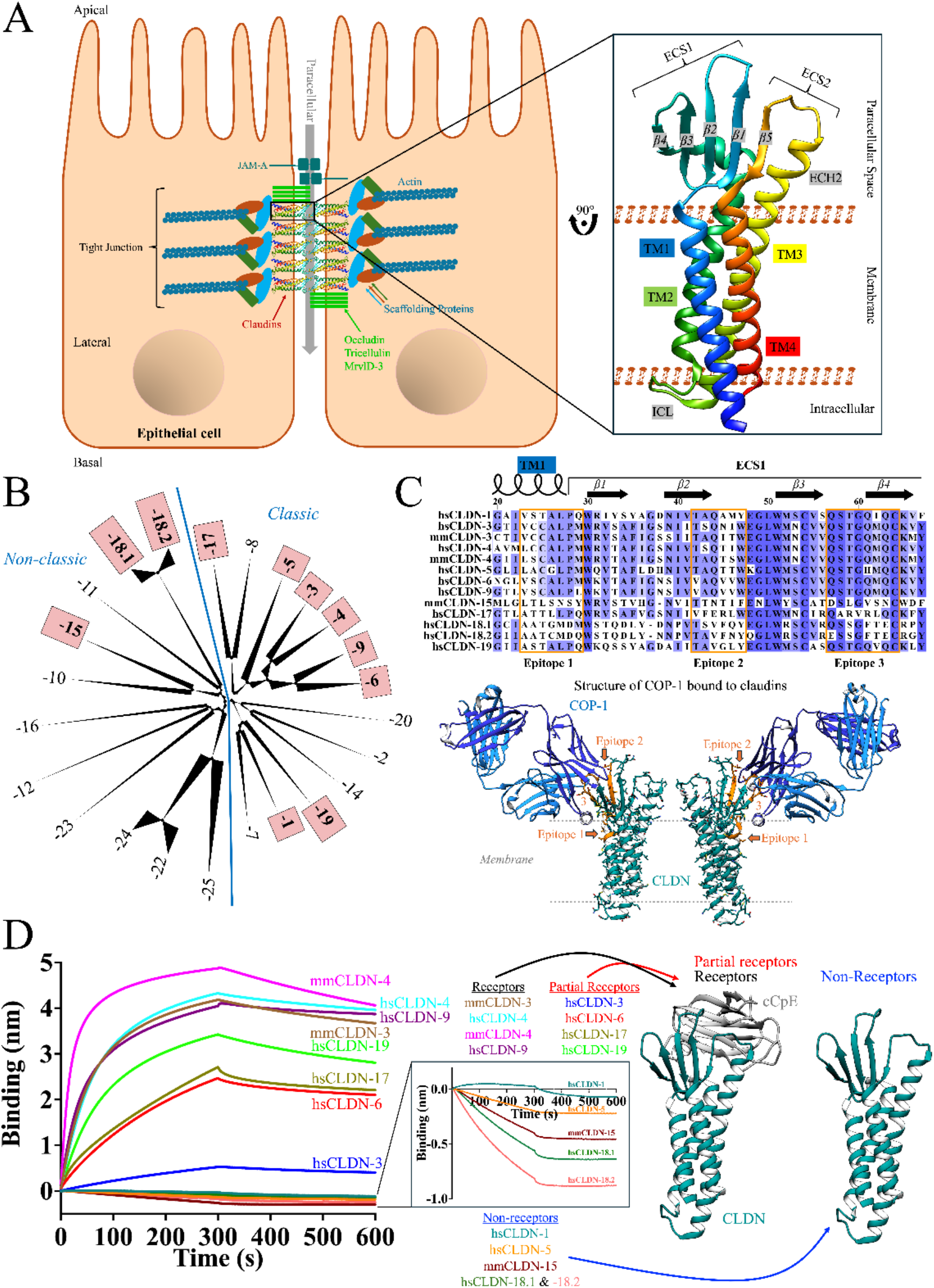
Sequence, structure, and function classification of claudins. Model of an epithelial bicellular tight junction with zoom-in depicting the 3D structures of claudins. Claudins are colored N-(blue) to C-terminus (red) and domains of interest are labeled as follows: transmembrane (TM) domain, extracellular segment (ECS), intracellular loop (ICL), and extracellular helix (ECH). (**B**) Phylogenetic tree of the claudin family highlighting classic vs. non-classic subtypes and subtypes used in this study (pink box). (**C**) The three known COP-1 binding epitopes on claudins. Sequence alignment of the 13 claudins used in this study with epitopes highlighted (orange box). Also shown is the structure of COP-1 (blue) bound the hsCLDN-4 (cyan) with epitopes 1-3 shown (orange). (**D**) Binding of claudins to cCpE from BLI. Association and dissociation phases are 300 seconds each. Zoom-in shows binding experiment with claudin non-receptors. Grouping of claudins into three categories based on cCpE receptor capacity from this experiment are shown in relation to structures of claudins alone or claudin/cCpE complexes.

Claudins with diverse tissue distributions and paracellular transport functions that are implicated in a wide array of diseases were chosen for this study. Specifically, claudin-1 is expressed in skin and maintains the epidermal paracellular barrier and is also a receptor for hepatitis C viral (HCV) entry in the liver (*6, 7*). Claudin-5 is the major regulator of the blood-brain barrier, which forms the protective diffusion layer that limits molecular exchange from the blood to the brain microenvironment and is thus being targeted by claudin-derived peptides, synthetic compounds, monoclonal antibodies and mutant *Clostridium perfringens* (CpE) enterotoxin to enable drug delivery into the brain (*8, 9*). Claudins -3 and -4 are highly expressed in the gastrointestinal tract, are barrier-forming claudins, and are receptors for *Clostridium perfringens* enterotoxin, a common cause of food poisoning (*10, 11*). Due to CpE interactions with claudins, it has been used to study claudin structure/claudin-receptor interactions and modified to develop mutants and peptidomimetics that modulate tight junctions for drug delivery or target cancers where claudins are overexpressed (*12*–*15*). Claudin-6 is expressed during embryonic tissue development where it regulates renal electrolyte homeostasis (*16*). Although it is not present in normal adult tissues, liver, ovarian, endometrial, testicular and esophageal cancers overexpress claudin-6, making it a target for solid tumor immunotherapies (*17*). Claudin-9 is essential for hearing by tuning sodium and potassium ion permeability ratios in subapical tight junctions of sensory hair cells in the inner ear (*18*). It was identified as a co-entry receptor, with claudin -1 and -6, for HCV infection (*7*). In the kidney, claudins -15, -17, and -19 form diverse ion-selective channels that regulate magnesium and calcium reabsorption, the latter through hetero-oligomeric interactions with claudin-16 (*19*– *21*). Claudin-18 splice variant 18.1 is highly expressed in lung alveolar epithelia where it regulates the alveolar fluid clearance, serves as the primary airway epithelial barrier to aero antigens and functions as a tumor suppressor attenuating malignant properties including cell proliferation, migration and invasion (*22*–*24*). Claudin-18.2 is normally expressed in differentiated epithelial cells of the gastric mucosa where it blocks paracellular gastric acid leakage from the gastric lumen into the submucosal space (*25*). It is also observed to be abnormally activated in pancreatic, esophageal, ovarian and lung tumors (*26*). Clearly, the vast yet unique functionalities of individual claudin subtypes play important roles in epithelial and endothelial tissue homeostasis while dysregulation or mis-assembly of tight junctions is a hallmark of numerous disease states.

Due to their roles in formation and maintenance of the tight junction barrier, claudins are attractive targets for the development of claudin-binding molecules, which hold potential to alter tight junction permeability to deliver drugs or to treat diseases linked to tight junction mis- or dis-assembly (*27*– *29*). To be effective, however, such molecules need to bind the paracellularly-exposed surfaces of claudins. We have described previously that targeting of claudins is challenging due to their small ∼25 kDa masses and having half of their mass being buried within membranes or disordered as termini (*30*). There is thus only ∼40% (10 kDa) worth of “targetable” extracellular surface area for antibodies or drugs to bind. The C-terminal domain of CpE (cCpE) is known to bind the extracellular domain of claudins and is known to modulate tight junction barrier permeability (*31*–*33*). However, cCpE only binds with <100 nM affinity to a subset of claudin subtypes and has been shown to be most effective at modulating epithelial barriers after basolateral but not apical delivery (*34, 35*). Thus, cCpE is not generalizable and may not be an optimal tight junction modulator. New claudin-binding molecules are needed to fill this gap in therapeutic potential.

Recently, we described a synthetic antibody fragment (sFab) termed COP-1 that was developed to target human claudin-4 (hsCLDN-4) (*30*). Structural analysis of COP-1 bound to hsCLDN-4 revealed that it binds the extracellular domain of hsCLDN-4 in a region different than cCpE and even accesses its transmembrane domain. Although COP-1 bound hsCLDN-4 best, we showed that it bound homologous claudins with high affinity too. This raised the possibility that COP-1 with its unique binding mode might bind other claudins. Here, we conduct detailed biophysical characterization of COP-1 binding to a diverse selection of 10 claudin subtypes from both humans and mice, which reveals that COP-1 binds with <100 nM affinity and with an identical mechanism to all claudins tested. Further, using a tissue system that models small intestinal epithelium and endogenously expresses at least six claudin subtypes, we show that COP-1 reversibly opens paracellular barriers. COP-1 is a newly identified claudin binding molecule distinct from cCpE whose pan-claudin-binding ability holds potential to increase tight junction permeability broadly across different tissues with diverse claudin expression patterns and barrier properties.

## Results

### Sequence and structure analysis of claudins used in this study

Claudin 3D structure has been elucidated previously and consists of four transmembrane (TM) domains, two extracellular segments (ECS) that span paracellular space, and intracellular N- and C-termini (**Fig. 1A**) (*30, 36*–*41*). The 27 human claudins were sequence aligned to reveal homology within the family (**Fig. S1** and **Table S1**). Phylogenetic analysis showed that claudins cluster into two distinct previously known groups termed classic and non-classic claudins (**Fig. 1B**) (*42*). Thirteen unique claudins, 10 from humans and three from mouse, representing ten subtypes were focused on for further study. These included human claudins -1, -3, -4, -5, -6, -9, -17, -18.1, -18.2, and -19 (hsCLDN-x); and murine claudins -3, -4, and -15 (mmCLDN-x). The selected claudins span sequence diversity, represent classic and non-classic subtypes, as well as receptors and non-receptors of CpE (*12, 40*–*43*).

From the structure of COP-1 bound to hsCLDN-4 three epitopes on claudins were identified that direct COP-1 binding (*30*). Alignment of residues spanning the three epitopes revealed that sequence conservation in these epitopes range from 19-91% in the 13 claudins of focus compared to hsCLDN-4—with hsCLDN-18.2 being most divergent and hsCLDN-3 being least divergent (**Fig. 1C**). Specifically, in epitope 1 we observe 0-89% (avg. 52%); in epitope 2, 17-83% (avg. 42%); and in epitope 3, 29-100% (avg. 74%) sequence identity. We have shown that mutants to claudin residues within these epitopes can decrease or increase COP-1s affinity to verify their involvement in binding (*30*). Having discerned how homologous or diverse each subtype was to hsCLDN-4, we embarked on biophysical analyses to discern how sequence divergence in COP-1 epitopes effect binding.

### Establishing the cCpE receptor capacities of claudins

We cloned, recombinantly expressed, then biochemically purified the 13 claudins using established methods (*40, 41*). First, because several claudins had not been isolated or functionally characterized before *in vitro*, we measured each claudin’s ability to bind cCpE, a known claudin-binding protein, using bio-layer interferometry (BLI). We found that claudins could be grouped into three classes based on estimated equilibrium dissociation constants (K_Ds_): those that bound cCpE with <20 nM affinity (claudins -3, -4, and -9), those that bound with 20-300 nM affinity (claudins -3, -6, -17, and -19), and those that did not bind cCpE (claudins -1, -5, 15, and -18) (**Fig. 1D** and **Table S2**). Interestingly, the human and murine orthologs of claudin-3 fit in two distinct classes. These results showed that the 13 claudins could be classified as cCpE receptors, partial receptors, and non-receptors, and agreed with binding analyses conducted previously (*30, 37, 40, 41, 44*).

Since 500 nM claudin was used to scout binding, we validated that non-receptors did not bind cCpE by increasing claudin concentrations to 2,000 nM. Here, claudins -5, -15, and -18 did not bind cCpE while hsCLDN-1 bound cCpE poorly, which agreed with its low binding capacity reported before (**Fig. 1D, inset**) (*41*). Additionally, we quantified cCpE binding to claudins -6, -17, and -19 because complete BLI measurements had not been reported for these subtypes. We measured K_Ds_ of 51.4 nM, 27.0 nM, and 27.5 nM for hsCLDNs -6, -17, and -19, respectively (**Fig. S2** and **Table S2**). In full, these results confirmed or established the cCpE receptor capacities of claudins, that homologous subtypes can bind cCpE with varied affinity owing to minute sequence alterations in key regions, and that the claudins we prepared were functional. This information was critical to benchmark against COP-1 binding studies.

### Quantification of COP-1 binding to claudins

To determine COP-1s claudin-binding ability, we again performed a BLI single-point analysis. We found that COP-1 bound all 13 claudins with <100 nM affinity that ranged from 59.0 to 99.6 nM with varied rates of association but similar rates of dissociation (**Fig. 2A** and **Table 1**). We then tested COP-1 binding to claudins in the presence of cCpE. This was done to determine if the two proteins had unique or overlapping binding sites and to determine how cCpE binding could alter COP-1. Again, we found that COP-1 bound all claudins (or claudin/cCpE complexes) and generally that cCpE receptor claudins (−3, -4, -6, -9, -17, and -19) yielded higher binding signal that non-receptor claudins (−1, -5, -15, -18), owing to the larger masses of claudin/cCpE complexes versus claudin alone detected by BLI (**Fig. 2B**). The cCpE alone did not bind COP-1. Analysis of the K_Ds_ revealed that COP-1’s affinity for mmCLDN-3, hsCLDN-4, and hsCLDN-18.1 increased when cCpE was present while for hsCLDN-17 and -19 it decreased—for all other claudins no significant change in COP-1 affinity occurred compared to cCpE alone (**Table 1**). COP-1 does not compete with cCpE and thus has a unique binding site yet is sensitive to claudin structural changes induced by cCpE.

**Table 1.** Single-point analysis of COP-1 binding to claudins.

| Subtype | K<sub>D</sub> (nM) |  | K<sub>D</sub> (nM) |
| --- | --- | --- | --- |
| hsCLDN-1 | 76.3 ± 1.5 | +cCpE | 73.2 ± 1.5 |
| hsCLDN-3 | 64.2 ± 1.1 |  | 58.5 ± 1.2 |
| mCLDN3 | 90.5 ± 1.2 |  | 44.0 ± 0.9 |
| hCLDN4 | 72.8 ± 0.9 |  | 42.4 ± 0.9 |
| mCLDN4 | 99.6 ± 1.2 |  | 86.2 ± 0.8 |
| hCLDN5 | 72.0 ± 1.1 |  | 65.5 ± 1.2 |
| hCLDN6 | 59.0 ± 0.9 |  | 59.2 ± 1.0 |
| hCLDN9 | 69.9 ± 1.2 |  | 74.8 ± 1.0 |
| mCLDN15 | 63.3 ± 1.3 |  | 61.2 ± 1.4 |
| hCLDN17 | 65.5 ± 0.9 |  | 86.4 ± 1.3 |
| hCLDN18.1 | 83.4 ± 1.0 |  | 56.7 ± 1.4 |
| hCLDN18.2 | 62.8 ± 1.1 |  | 55.6 ± 1.3 |
| hCLDN19 | 62.4 ± 1.1 |  | 93.9 ± 1.4 |

**Fig. 2.**
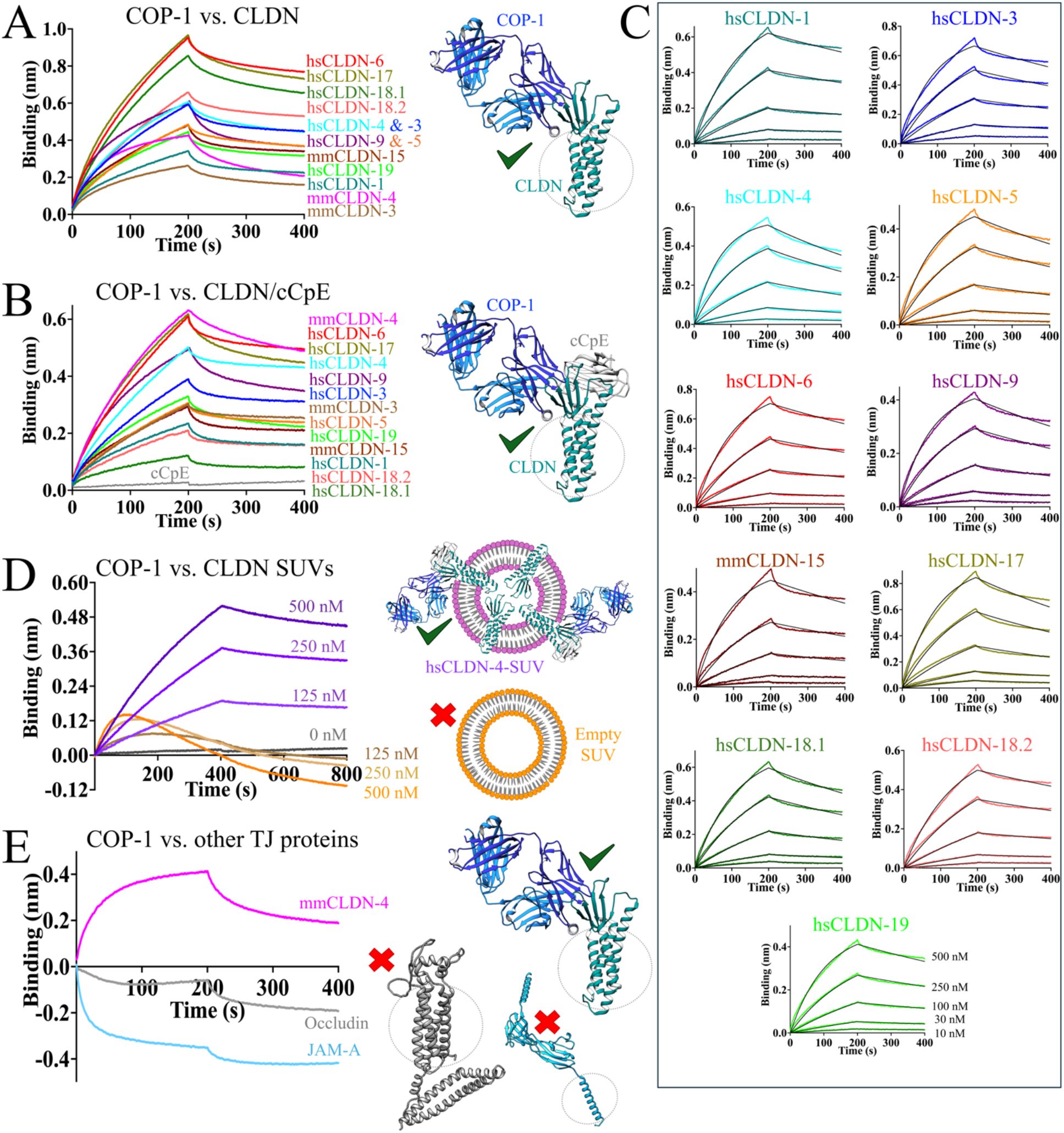
Biophysical characterization of COP-1 to claudins in various formats. (**A**) Single-concentration point binding assessment (500 nM) of COP-1 to 13 claudin subtypes shows binding signals for all. **Table 1** shows associated K_Ds_ values. (**B**) Full multi-concentration point (0-500 nM) analyses of COP-1 binding to 11 representative claudin subtypes. **Table 2** shows associated kinetic rates and K_Ds_ values. (**C**) Single-concentration point binding analysis (500 nM) of COP-1 to 13 claudin subtypes in the presence of cCpE. **Table 1** shows associated K_Ds_ values. (**D**) Multi-concentration point (0-500 nM) analyses of COP-1 binding to SUVs loaded (purple) or unloaded (orange) with hsCLDN-4. **Table 2** shows associated kinetic rates and K_Ds_ values for hsCLDN-4-SUVs. (**E**) Single-concentration point binding analysis (500 nM) of COP-1 binding to a control claudin (mmCLDN-4, pink) and human occludin (grey) and JAM-A (lt. blue) in detergent.

To validate that COP-1 binds claudins specifically, we tested a homologous sFab discovered simultaneously with COP-1 called COP-2 for its ability to bind claudins (*45*). COP-2 was known to bind cCpE and hsCLDN-4/cCpE complexes but not hsCLDN-4 alone (*45*). Single-point BLI using COP-2 showed that it did not bind claudins in absence of cCpE (**Fig. S3A**). However, when cCpE is added to claudins, COP-2 was found to bind to free cCpE in non-receptor/cCpE mixtures or to cCpE in receptor/cCpE complexes (**Fig. S3B**). These experiments validated that COP-1 is unique in its claudin-binding ability due to sequence changes in its complementarity-determining regions (CDRs) compared to other sFabs that form nearly identical tertiary structures.

We next quantified COP-1 binding to all 10 claudin subtypes more extensively. We measured K_Ds_ of 61.3 nM for hsCLDN-1, 57.9 nM for hsCLDN-3, 70.6 nM for hsCLDN-4, 75.8 nM for hsCLDN-5, 59.5 nM for hsCLDN-6, 66.0 nM for hsCLDN-9, 59.0 nM for mmCLDN-15, 80.2 nM for hsCLDN-17, 92.4 nM for hsCLDN-18.1, 66.4 nM for hsCLDN-18.2, and 62.2 nM for hsCLDN-19 (**Fig. 2C**). This showed that COP-1 bound claudins with a range of 58-92 nM (**Table 2**). Using the dissociation rate (k_off_), we calculated the half-life (t_1/2_) of claudin/COP-1 complexes, which revealed that the t_1/2_ of claudin/COP-1 complexes were between 8-13 minutes. We tested whether claudin/COP-1 complexes could be retained in solution using analytical size-exclusion chromatography (SEC) to validate this finding. By comparing the elution times of claudins alone (**Fig. S4A**) versus claudins mixed with COP-1 and a sFab-specific nanobody (**Fig. S4B**), we found that COP-1s presence decreased average elution times from 5.6 to 5.1 minutes (**Fig. S4C**) (*46*). This verified the formation of larger complexes at 5.1 minutes. COP-1 alone eluted at 7.2 minutes. These results suggested that the claudin/COP-1 complex is stable *in vitro* but is relatively short-lived, prompting us to determine whether COP-1 binds differently to claudins in membranes—a prerequisite for therapeutic applications.

**Table 2.** Full quantification of COP-1 binding to claudins.

| Subtype | Mimetic | K<sub>D</sub> (nM) | k<sub>on</sub> (1/Ms) | k<sub>off</sub> (1/s) | t<sub>1/2</sub> (min) | Epitope ID (%) |
| --- | --- | --- | --- | --- | --- | --- |
| hCLDN1 | Detergent | 61.3 ± 0.5 | 1.5x10<sup>4</sup> ± 0.7x10<sup>2</sup> | 0.9x10<sup>-3</sup> ± 5.4x10<sup>-6</sup> | 12.8 | 57 |
| hCLDN3 |  | 57.9 ± 0.5 | 2.1x10<sup>4</sup> ± 1.1x10<sup>2</sup> | 1.2x10<sup>-3</sup> ± 7.3x10<sup>-6</sup> | 9.6 | 91 |
| hsCLDN-4 |  | 70.6 ± 0.4 | 2.6x10<sup>4</sup> ± 1.2x10<sup>2</sup> | 1.8x10<sup>-3</sup> ± 7.1x10<sup>-6</sup> | 6.4 | 100 |
| hCLDN5 |  | 75.8 ± 0.5 | 1.9x10<sup>4</sup> ± 0.9x10<sup>2</sup> | 1.5x10<sup>-3</sup> ± 6.4x10<sup>-6</sup> | 7.7 | 76 |
| hCLDN6 |  | 59.5 ± 0.4 | 1.8x10<sup>4</sup> ± 0.9x10<sup>2</sup> | 1.1x10<sup>-3</sup> ± 6.0x10<sup>-6</sup> | 10.5 | 71 |
| hCLDN9 |  | 66.0 ± 0.4 | 2.2x10<sup>4</sup> ± 0.9x10<sup>2</sup> | 1.5x10<sup>-3</sup> ± 5.9x10<sup>-6</sup> | 7.7 | 67 |
| mmCLDN-15 |  | 59.0 ± 0.5 | 2.1x10<sup>4</sup> ± 1.2x10<sup>2</sup> | 1.2x10<sup>-3</sup> ± 8.2x10<sup>-6</sup> | 9.6 | 33 |
| hCLDN17 |  | 80.2 ± 0.6 | 1.9x10<sup>4</sup> ± 1.1x10<sup>2</sup> | 1.5x10<sup>-3</sup> ± 7.8x10<sup>-6</sup> | 7.7 | 29 |
| hCLDN18.1 |  | 92.4 ± 0.5 | 1.6x10<sup>4</sup> ± 0.7x10<sup>2</sup> | 1.4x10<sup>-3</sup> ± 4.9x10<sup>-6</sup> | 8.3 | 33 |
| hCLDN18.2 |  | 66.4 ± 0.5 | 1.3x10<sup>4</sup> ± 0.7x10<sup>2</sup> | 0.9x10<sup>-3</sup> ± 5.2x10<sup>-6</sup> | 12.8 | 19 |
| hsCLDN-19 |  | 62.2 ± 0.4 | 1.7x10<sup>4</sup> ± 0.7x10<sup>2</sup> | 1.1x10<sup>-3</sup> ± 5.3x10<sup>-6</sup> | 10.5 | 57 |
| hsCLDN-4 | SUV | 105.7 ± 0.3 | 0.4x10<sup>4</sup> ± 0.1x10<sup>2</sup> | 0.4x10<sup>-3</sup> ± 0.7x10<sup>-6</sup> | 28.9 | 100 |

### COP-1 binds claudin-4 in membranes

Because claudins are membrane proteins and can oligomerize within membranes, we tested if COP-1 bound claudins in membranes in these altered states as a predictor of whether they could be useful *in vivo*. Indication that it might came from our finding that COP-1 bound hsCLDN-4 reconstituted in lipid nanoparticles called nanodiscs, which possesses a small lipid bilayer but is small enough to only contain a claudin monomer (*47, 48*). We reconstituted hsCLDN-4 in small unilamellar vesicles (SUVs) composed of the lipid diphytanoylphosphatidylcholine (DPhPC), which has been shown to produce functional claudin-4, and compared COP-1 binding to empty DPhPC SUVs (*49*). We found that empty SUVs gave a non-sensical and non-concentration dependent binding signal whereas hsCLDN-4-loaded SUVs gave concentration dependent binding signals with robust kinetic rates (**Fig. 2D**). We calculated a K_D_ and t_1/2_ from the rates, which showed that COP-1 bound hsCLDN-4 in SUVs with 105.7 nM affinity and that the complex half-life approached 30 minutes (**Table 2**). The rates of COP-1 association (k_on_) and dissociation (k_off_) for hsCLDN-4 were >4-fold slower in SUVs compared to detergent. This finding showed that COP-1 binds claudins in membranes in physiologically relevant states and that the complex is long-lived, piquing our interest to test its effect on claudins residing in epithelial cells.

### COP-1 does not bind other tight junction membrane proteins

Before deciphering COP-1s binding in model tissues we established whether COP-1 could bind other membrane proteins with paracellularly-exposed domains found at tight junctions to rule out off-target effects. Occludin increases paracellular barrier function by driving the formation and stabilization of tight junction branching points (*50, 51*). Junctional adhesion molecule-A (JAM-A, also known as the F11 receptor) regulates membrane apposition and mediates intercellular signaling events at tight junctions while also serving as the large molecule barrier protein (*52*–*54*). We expressed and purified human occludin and JAM-A and found that they did not bind COP-1, which was verified by comparing to mmCLDN-4 (**Fig. 2E**). This result confirmed that COP-1 binds selectively to claudins and that any potential effects on barrier function observed *in vivo* were likely to be a result of this selectivity and not binding to other paracellularly exposed tight junction membrane proteins.

### COP-1 alters tight junction barrier permeability in an intestinal tissue model

We cultured Caco-2 cells and measured transepithelial electrical resistance (TEER) to evaluate the effect of COP-1 on tight junction barrier permeability. Confluent Caco-2 monolayers were treated at their apical and/or basolateral compartments with 500 nM COP-1 and various control proteins that included 500 nM cCpE, 200 nM CpE, and 500 nM COP-2. CpE is known to kill intestinal epithelial cells while cCpE has been shown to modulate tight junction barrier permeability in tissue model monolayers (*34, 55, 56*). TEER was measured in transwell plates before the addition of test proteins and 24 hours after treatment with a STX chopstick electrode. TEER after treatment was compared to pre-treatment for all proteins. Results show that COP-1-treated monolayers exhibited decreases in TEER of 19.6% and 37.3% after apical and basolateral delivery, respectively (**Figs. 3A** and **S5A**). The control protein cCpE decreased TEER by 12.7% when applied apically and 62.8% when applied basolaterally, consistent with previous findings (*34, 57*). Further, results of apical delivery of COP-2 and CpE showed that COP-2 had no effect while CpE reduced TEER to ∼0%, indicating obliteration of tight junction barriers after CpE-induced cytotoxicity (*55*). These results established that COP-1 modulates paracellular permeability on the order of cCpE.

**Fig. 3.**
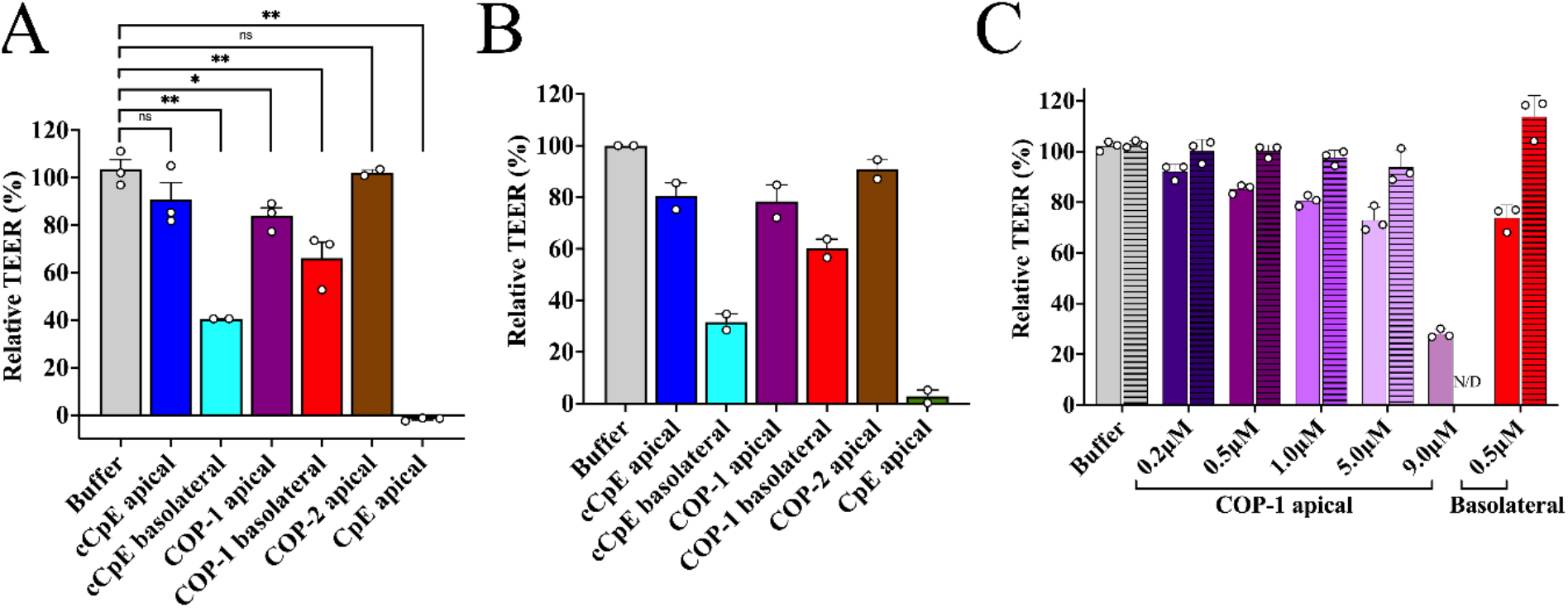
Effect of COP-1 on tight junction barrier integrity in a model for intestinal epithelium. (**A**) Plot of relative TEER measurements using an STX electrode after delivery of various proteins to the apical or basolateral compartments of Caco-2 monolayers. Proteins include cCpE, COP-1, COP-2, and CpE. TEER was measured n=2 for cCpE basolateral and COP-2 apical treatments, and measured n = 3 for buffer, COP-1 apical and basolateral, cCpE and CpE apical treatments. (**B**) Plot of relative TEER measurements using an Ussing chamber from the same monolayers used in **A**. TEER was measured n = 2 for all concentrations. (**C**) COP-1 concentration-dependent decrease in TEER. A concentration range (0-9000 nM) of COP-1 was added to apical compartments and single concentration (500 nM) added to basolateral compartments of Caco-2 cells and TEER was measured using a STX electrode, n = 3. The recovery of barrier function was measured after removal of COP-1. Solid bars represent data from 24 hours while patterned bars represent data from 48 hours (that is 24 hours after COP-1 containing media is exchanged with fresh medium), n = 3. The 48-hour time-point for 9 µM COP-1 apical measurement was not determined (N/D). All data are represented as mean ± SEM of two or three independent measurements. * represents p< 0.05 and ** represents p< 0.001 in TEER from treated cells compared to buffer alone.

TEER was additionally measured 24 hours after treatment with an Ussing chamber to validate the STX recordings. Here, Caco-2 monolayers were mounted in Ussing chambers in Ringer buffer solution and TEER was measured. We found that TEER levels decreased by 21.6% and 39.8% after COP-1 delivery to apical and basolateral compartments, respectively (**Fig. 3B**). Caco-2 cells treated with cCpE decreased TEER by 19.6% and 68.4% after identical delivery. Apical delivery of COP-2 treatment yielded no significant changes to TEER while CpE decreased TEER by 97.2%. These findings agreed with the STX electrode recordings and confirmed that COP-1 modulated tight junction barrier permeability in a manner and magnitude similar to cCpE.

We next determined whether Caco-2 tight junctions could be affected by COP-1 in a concentration-dependent manner, which was important to ascertain for therapeutic use. We found after COP-1 application to apical compartments that TEER decreased by 10.0% at 200 nM, 16.9% at 500 nM, 21.5% at 1000 nM, 29.1% at 5000 nM, and 71.0% at 9000 nM as measured using a STX electrode after 24 hours treatment compared to buffer alone (**Figs. 3C** and **S5B**). We then determined if tight junction permeability was reversible by removing COP-1 containing medium, exchanging it for fresh medium, then measuring TEER 24 hours after exchange. We found that TEER increased back to pretreatment levels for monolayers treated with <1000 nM COP-1 but not for monolayers treated with >1000 nM COP-1 (**Fig. 3C**). We also found in monolayers treated with 500 nM COP- 1 at their apical or basolateral compartments, that TEER recovered more completely in basolaterally-treated cells. Altogether, these results indicated that COP-1 increases paracellular permeability of a model epithelium in a concentration-dependent and reversible manner, and that this modulating effect may be prolonged upon apical delivery.

### Structural basis of COP-1 binding to claudins

Lastly, we attempted to elucidate the structural basis of COP-1’s claudin-binding ability and to assess whether COP-1 could enable structure determination of claudins. Because claudins are dynamic and low molecular weight proteins, they are recalcitrant to structural determination by cryogenic electron microscopy (cryo-EM). The sFabs COP-1 and -2 were developed to act as fiducial marks to enable structures by cryo-EM (*30, 45*). We incubated COP-1 and the sFab nanobody (Nb) with claudins from each of the three cCpE receptor classes and isolated claudin/COP-1/Nb complexes by SEC (**Fig. 4A**). The complexes were vitrified on grids and imaged using either 200 kV or 300 kV cryo-EM microscopes. Processing of the cryo-EM data yielded 2D classifications representative of claudin/COP-1/Nb complexes where the sFab/Nb is observed bound to a circular mass (**Fig. 4B**). Interestingly, the quality of 2D classes and presence of secondary structural elements was related to the cCpE receptor capacity of the claudin analyzed, where receptors yielded better classes than partial receptors and non-receptors were of lowest quality. This finding was not a result of number of particles processed or whether higher energy microscope were used. 3D reconstructions yielded maps of variable quality and of moderate resolution (4-9 Å) where COP-1/Nb were well-resolved but the claudins were not (**Fig. S6**). The detergent belt encasing claudins appears to dominate the signal and masks the claudin, making them largely unresolvable in the maps. Because of the high confidence we had in COP-1/Nb placement we modeled the structure of hsCLDN-4/COP-1/Nb bound to cCpE (PDB ID: 8u4v) into the maps, which showed that COP-1 binds all claudins tested in an approximately identical manner (**Fig. 4C**). Although COP-1 was not able to yield high resolution structures of claudins by cryo-EM, these results verified our biochemical and biophysical findings and provided validating structural information for COP-1s claudin binding mechanism.

**Fig. 4.**
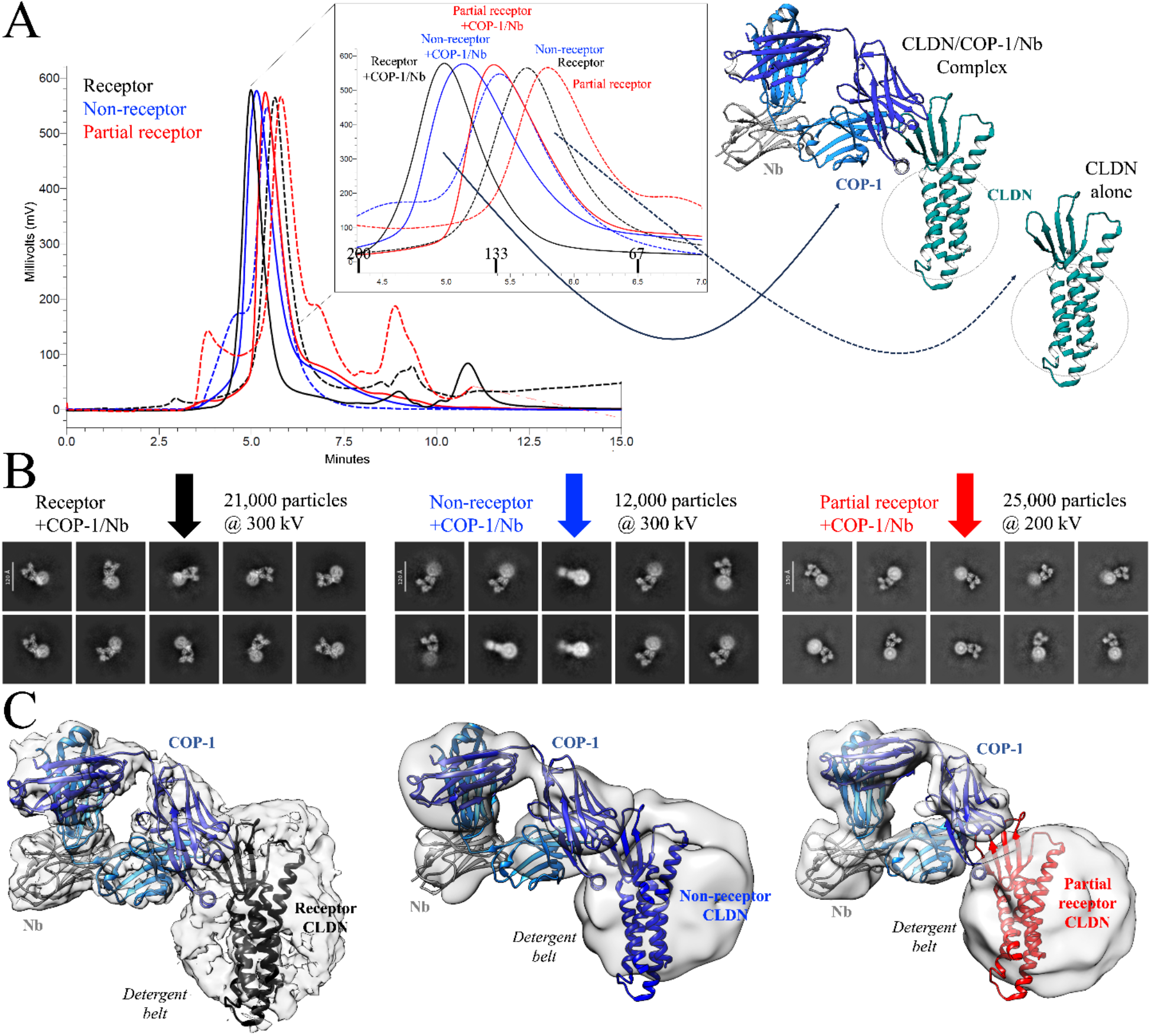
Structures of COP-1 bound to claudins by cryo-EM. (**A**) SEC chromatograms showing the elution times of claudins alone (dashed lines) versus claudin/COP-1/Nb complexes (solid lines). Structural models of the expected complexes are shown based on PDB ID 8u4v where claudins (teal) and COP-1 (blue) are depicted as cartoons. (**B**) 2D classifications and (**C**) 3D reconstructions from cryo-EM of claudin/COP-1/Nb complexes from the three cCpE receptor classes (receptors, black; non-receptors, blue; partial receptors, red). Note that the quality of 2D classes and final maps are related to claudin cCpE receptor classification, indicating that COP-1 stabilizes receptor claudins better than partial or non-receptor claudins.

## Discussion

We developed sFabs against cCpE and hsCLDN-4 to enable structure determination by cryo-EM (*30, 45*). Upon structural and biophysical characterization of the hsCLDN-4-binding sFab, COP-1, we discovered cross-reactivity with homologous claudins and that the epitopes that COP-1 used to bind hsCLDN-4 had sequence identity across the claudin family (**Fig. 1C**). This initially allowed us to predict that COP-1 could also bind claudins -3, -5, -6, and -9 (*30*). Here, we isolated 13 distinct claudins representing 10 subtypes and tested COP-1s claudin-binding ability, benchmarking it against the well-characterized claudin binding protein cCpEs. Applying our findings, we can classify the 13 claudins tested into three categories based on their affinity for cCpE, which include receptors (mmCLDN-3, hsCLDN-4, mmCLDN-4, hsCLDN-9), partial receptors (hsCLDN-3, hsCLDN-6, hsCLDN-17, and hsCLDN-19), and non-receptors (hsCLDN-1, hsCLDN-5, mmCLDN-15, and hsCLDN-18.1 and -18.2). This classification stems from the ability of claudins to bind cCpE at 500 nM, which is slightly above the pathophysiological concentration maximum of 350 nM (*58*). Coupled with the sequence diversity within claudins that define classic and non-classic subtypes, cCpE receptor capacity becomes a useful characteristic to interpret the structure and function of claudins. In this study, it becomes important to benchmark COP-1 function and potential application against cCpE. Additionally, and more importantly, we establish here that COP-1 is a high-affinity (<100 nM) and pan-claudin binding molecule with a novel binding mode and binding site that encompasses the transmembrane and extracellular domains of claudins, which is distinct from cCpE’s. In full, our binding analyses demonstrate that COP-1 binds uniquely and more broadly to claudins compared to cCpE, allowing us to surmise that COP-1 could be used generally to alter tight junction barrier function. Critically and unlike cCpE, which is most useful to target tissues that highly express CpE receptor claudins, COP-1 binds classic and non-classic and CpE receptor and non-receptor claudins equally, offering the possibility that it could open tight junctions as cCpE does but in non-tissue-specific ways.

Our biophysical results show that COP-1 binding to claudins is not largely altered in either a positive or negative way in the presence of cCpE, indicating that its binding epitopes are structurally similar when bound or unbound to cCpE (**Table 1**). This finding makes sense because cCpE binds the palm region of the claudin hand, so COP-1 freely binds epitopes that reside on the back of the claudin hand (**Fig. 2B**). This led us to speculate that COP-1 could bind claudins in any environment if these epitopes are accessible. We validated this theory by showing that COP-1 binds hsCLDN-4 in SUVs but does not bind claudin-less SUVs (**Fig. 2D**). This finding is significant as the structures of claudins and their oligomeric states in membranes are unknown and indicates that claudin structure in detergents approximates those in membranes and that COP-1 binds claudins directly. We provide further evidence that COP-1 binding is selective to claudins with our data that shows COP-1 is unable to bind occludin or JAM-A, two other tight junction membrane proteins that are structurally unique from claudins but which also have paracellular exposed domains (**Fig. 2E**). Notably, COP-1 distinguishes claudins from occludin, which have a similar four transmembrane domain architecture but larger ECS (*50*). Our findings verify that COP-1 is a selective claudin-binding molecule and that COP-1 is optimized to bind the claudin fold. We hypothesize and would be interested to test whether COP-1 binds other claudin-like proteins in the greater PMP22/EMP/MP20/claudin/voltage-gated calcium channel γ subunit family (pfam00822 superfamily). These integral membrane proteins perform a plethora of cellular functions at cell/cell contacts and beyond in invertebrates and vertebrates, and thus COP-1 could potentially modulate their functions in protein-specific ways—expanding its applicability (*59*).

COP-1s claudin-specific and paracellular domain binding, like cCpE, makes it an ideal molecule to alter normal tight junction function so we tested this in a model intestinal epithelium. COP-1 disrupts tight junction barrier function in Caco-2 monolayers after both apical and basolateral application and this disruption is reversible (**Fig. 3**). Apical delivery of COP-1 also decreases TEER in a concentration-dependent manner and once removed, tight junction barrier function is restored, although the magnitude of recovery is also concentration-dependent. Interestingly, barrier recovery appears to be more complete if COP-1 is delivered to basolateral versus apical compartments. Compared to the known tight junction disruptor cCpE (*34, 35*), COP-1 decreases barrier integrity equivalently when applied apically; whereas basolateral treatment shows cCpE is more effective at barrier disruption. In full, our data suggests that COP-1 and cCpE open paracellular barriers to similar degrees using similar mechanisms—the details of which can be elucidated by considering how claudins integrate into tight junctions and the proportion of cCpE receptor claudins in a given tissue.

Van Itallie *et al*. showed that new claudins, those yet to polymerize, are integrated into tight junction strands through basolateral pools in order to keep apical barriers intact (*60*). Old claudins are subsequently removed from apical strands in unknown oligomeric states to then perform other functions or to be degraded. Caco-2 cells express claudins -1, -2, -3, -4, -7 and -15 (*57, 61*). These represent classic and non-classic and all three classes of cCpE receptor (**Figs. 1B** and **1D**). Basolateral delivery of cCpE will readily bind non-polymerized claudins -3 and -4 and prevent their integration into tight junction strands, affecting barrier function. This “sequestering” mechanism first described by Sonoda *et. al* is based on their finding that basolateral but not apical delivery of cCpE to MDCK monolayers decreased TEER posits that cCpE disrupts the equilibrium between non-polymerized and polymerized claudins (*34*). The decreases to Caco-2 monolayer TEER we find after basolateral delivery of both cCpE and COP-1 agrees with a sequestering mechanism— with the difference being that cCpE binds only receptor claudins whereas COP-1 binds all claudins—yet both sequester claudins from integration into tight junctions. We speculate that the greater effect on TEER by basolateral cCpE can be explained by several factors. One, intestinal epithelium and Caco-2 cells are rich in cCpE receptors, so cCpE can have a greater impact on barrier function in such tissues (*57, 61*). Two, cCpEs higher affinity binding and longer complex half-life for claudins would sequester them better and longer from polymerizing compared to COP-1 (**Table S2**). Three, cCpE binding is known to alter the structures of claudins while COP-1 adapts naturally to their structures (*30, 37, 38, 40, 62*). And four, the COP-1 epitope may not be as accessible in polymerized or non-polymerized claudins as cCpEs binding surface—more on this to follow. These factors, coupled with the integration of claudins into tight junctions from basolateral compartments, explain why cCpE is a better modulator of barrier function when applied basolaterally.

Apical delivery of cCpE and COP-1 decreased TEER equally while higher amounts of COP-1 decreased TEER further in concentration-dependent and reversible ways (**Fig. 3**) (*55*). These findings suggest that cCpE and COP-1 bind polymerized claudins within tight junctions. In the more developed tight junctions of apical epithelium, old claudins are removed from active strands but new ones are not synthesized from there [*56*]. Thus, cCpE or COP-1 binding to non-polymerized apical claudins would not alter the polymerized/non-polymerized equilibrium and not affect barrier integrity as greatly. This idea agrees with our results that apical treatments decrease TEER less that basolateral application and that TEER recovers more completely after basolateral treatment of COP-1 compared to apical. In order to affect the barrier integrity of apical epithelium, cCpE and COP-1 must bind polymerized claudins and subsequently alter their structures and the structure of tight junctions. This is more challenging because unlike non-polymerized claudins, polymerized claudins are proposed to associate in distinct ways that may shield cCpE and COP-1 binding surfaces. Our results suggest that cCpEs and COP-1s binding epitopes are at least partially accessible in claudin-polymerized tight junctions because these proteins decrease TEER when applied apically. These findings provide further evidence that COP-1 may be as potent of a tight junction modulator as cCpE but that it could have enhanced applicability as it does not rely on specific receptors.

Shrestha *et al*. found that 285 nM cCpE delivered apically to Caco-2 cells decreased TEER by 50% overnight while we found that 500 nM COP-1 decreased TEER by ∼22% (*55*). This suggests that although both proteins target claudins in tight junctions, that like basolateral treatment, cCpE is more effective. Again, this could be because Caco-2 cells are richer in receptor claudins and cCpE is better at binding and altering these subtypes than COP-1 is at doing so generally; or, that there are differences in subtype composition in apical versus basolateral strands, as has been suggested and shown (*58, 63*); or, that the COP-1 epitope is not as accessible as cCpEs binding surface in polymerized claudins. To understand how cCpE, COP-1, or any molecule alters claudin structure to affect apical tight junction barriers requires structures of polymerized claudins. Unfortunately, no experimental structures exist. However, using models of polymerized claudins we find that cCpE and COP-1 binding surfaces may indeed be differentially accessible (*5, 36, 64*– *66*). In straight lateral assemblies both cCpE and COP-1 surfaces are accessible (**Fig. S7A**) (*36*); in face-to-face dimers (X-1), only cCpEs surface is accessible (**Fig. S7B**) (*65*); in an alternate dimer (Cis-1), cCpEs surface is accessible but COP-1s is partially occluded (**Fig. S7C**) (*64*); and in a yet verified dimer that we generated, both protein’s surfaces are accessible (**Fig. S7D**). These models of polymerized claudins suggest that cCpEs binding surface is available in more claudin polymers than COP-1s epitopes. We hypothesize that unlike cCpE, COP-1 could be used to elucidate the structure and organization of polymerized claudins in different tissues as it can only bind effectively to distinct claudin polymers. Further validation of these models or determination of experimental structures of polymeric claudins are needed to test this idea and to develop better claudin-binding molecules.

In sum, COP-1 is a novel and specific pan-claudin-binding molecule capable of reversibly modulating paracellular permeability in tissues through targeted disruption of tight junctions. COP-1 has the potential for “tunability” because the apical route directly insults tight junctions and recovers slower that the basolateral route, which is more dynamic due to being the site of claudin synthesis and integration. Although generated unnaturally, COP-1 is nearly as effective at opening tight junction barriers in intestinal tissues as a highly evolved, natural, and subtype-selective tight junction modulator, cCpE. COP-1s pan-claudin-binding ability may enable it to outperform cCpE in barrier disruption of more diverse epithelia or endothelia. Explicitly, we surmise that COP-1s insult to tight junction barrier integrity will be greater than cCpEs in tissues lacking cCpE receptor claudins but rich in other subtypes because both modulators have overlapping mechanisms of barrier disruption but unique modes of binding. Thus, for modulating paracellular permeability, COP-1 may have increased potency while its claudin-specificity yields less off-target effects. As is, or with increased development, COP-1 or its derivatives could serve a wide-range of translational applications that include modulation of tight junctions, drug delivery through paracellular spaces, detection of claudin-rich cancers, or inhibition of inflammatory or oncogenic pathways.

## Materials and Methods

### Sequence Analysis of Claudins

All claudin sequences were obtained from UniProt database (*67*). We performed multiple sequence alignment and phylogenetic analysis with T-coffee on the EMBL-EBI webserver (*68, 69*). The aligned claudin sequences were viewed and colored according to sequence identity in Jalview (*70*). The secondary structural elements, as marked on the sequence alignment was mapped out based on human claudin-4 structure (PDB ID: 7kp4).

### Protein Expression and Purification

All claudins and cCpE were expressed and purified as previously described (*30, 40, 41*). All proteins contained a terminal poly-histidine tag with thrombin protease cleavage site. Proteins were expressed in insect cells, purified by immobilized metal affinity chromatography, then released from NiNTA resin after thrombin digestion. Protein purity and homogeneity was verified by SDS-PAGE and SEC, then concentrated stocks (1 mg/mL) were snap frozen in liquid N and stored at -80°C until use.

COP-1 and -2 with and without a C-terminal histidine tag were expressed in BL21 (DE3) cells using pRH2.2 plasmid encoding the gene for the protein as previously described (*30, 45*). Cells transformed with COP-1 plasmid were used to inoculate a 1 L overnight culture in TB media containing 100 µg/mL ampicillin. Cells were grown to and O.D. of 0.8 then induced with 1 mM IPTG. Cells were harvested by centrifugation after 4 hours at 37°C. Cells were lysed by sonication then lysate was incubated at 65°C for 30 minutes then rapidly chilled on ice for 15 minutes. 1 M NaCl and 0.01% n-dodecyl-β-D-maltoside (DDM, Anatrace) was added then lysate was ultracentrifuged at 8,200 xg for 30 minutes. Filtered supernatant was loaded onto a 5 mL protein L resin column (Cytiva) equilibrated in 20 mM Tris pH 7.4, 100 mM NaCl, 1% glycerol, and 0.01% DDM. COP-1 was eluted in 100 mM acetic acid then dialyzed in BLI buffer (20 mM Tris pH 7.4, 100 mM NaCl, 1% glycerol, 0.03% DDM). COP-1 was concentrated to 1 mg/mL and snap frozen in liquid N and stored at -80°C until use.

### Reconstitution of hsCLDN-4 in small unilamellar vesicles

35 mg of DPhPC was dried under nitrogen gas for 90 minutes then rehydrated in 2 mL of SUV buffer (25 mM Hepes 7.4, 150 mM NaCl, and 0.5 mM TCEP). The solution was tip sonicated for 4 minutes to make SUVs then filtered 10x through a 0.2 µm syringe filter (Pall). 140 µL of SUV buffer with 1% n-octyl-β-D-glucoside (OG) was used to make empty SUVs, while 70 µL of 2 mg/mL hsCLDN-4 in DDM was mixed with 70 µL of SUV buffer with 2% OG to make hsCLDN-4-SUVs. To 140 µL of both solutions, 340 µg of DPhPC SUVs and SUV buffer were added to 175 µL and incubated on the bench for 30 minutes. This solution was diluted 4-fold then placed in 3 mL Slide-a-lyzer 10 kDa MWCO cassette (Pierce) and dialyzed against 250 mL of SUV buffer in the presence of 2 g SM2-biobeads (Bio-Rad) overnight at 4°C. 140 µg of hsCLDN-4 in 875 µL yielded 0.16 mg/mL (6.95 µM) This stock was used for BLI analyses, despite only half of hsCLDN-4s may be oriented outside-out.

### Bio-layer Interferometry (BLI)

BLI was performed on an Octet R8 (Sartorius) at 25°C in 96-well black flat bottom plates (Greiner) using an acquisition rate of 5 Hz averaged by 20. Histidine-tagged proteins (500 nM cCpE, 100 nM COP-1, or 100 nM COP-2) in BLI buffer were immobilized on Ni-NTA Dip and Read biosensors then a kinetic experiment was performed that consisted of baseline (100 seconds), association (200-300 seconds), and dissociation (200-300 seconds) steps. Claudins in BLI buffer (0-2000 nM) or claudin/cCpE complexes (250 nM each) were present in association wells for most analyses. For binding of COP-1 to SUVs, SUV buffer was used and 100 nM COP-1 or 500 nM cCpE was tested against 0-500 nM hsCLDN-4-SUVs. The binding of hsCLDN-4-SUVs to immobilized cCpE verified that some claudin ECS were outside-out facing. An equal volume of empty SUVS were used, as these had no Abs_280nm_ due to not being loaded with a protein. All binding measurements were fit to a 1:1 binding model using BLitz Pro 1.3 Software then the kinetic data and fits were exported is .csv format and re-plotted using GraphPad Prism version 10.0.3.

### Size-exclusion Chromatography (SEC)

To assess COP-1 binding to claudins using SEC, 100 to 250 µg of claudin was added to 1 molar equivalent of COP-1 and incubated at 4°C overnight while nutating. 1.2 molar excess of the anti-sFab nanobody to COP-1 was added to the complex and incubated for 2 hours. These were 0.2 µm filtered and loaded onto a Superdex 200 increase (5/150) gl (Cytiva) column equilibrated in BLI buffer. The formation of complex between claudin and COP-1 was assessed by comparison of claudin alone elution times to claudin/COP-1/Nb elution time.

### Transepithelial Electrical Resistance (TEER)

Human colorectal adenocarcinoma cells (Caco-2) cells, obtained from the American Biological Culture Association, were a kind gift from the Duffey Lab, Department of Physiology and Biophysics, University of Buffalo. Cells were grown and maintained in DMEM, high glucose, GlutamMAX supplement, pyruvate (Gibco) supplemented with 10% Fetal Bovine Serum (FBS), 1x MEM Non-essential amino acid solution (Sigma Aldrich) and 1% penicillin-streptomycin. Caco-2 cells were seeded onto transwell plates (12mm pore, 0.4µm pore polycarbonate membrane, Corning Costar) and grown to confluency for at least 14 days at 37°C with 5% CO_2_ with a change of media every 48 hours. For the assay, confluent monolayers were treated with medium (no serum) containing 500 nM cCpE, 500 nM COP-1, 500 nM COP-2, 200 nM CpE or buffer for 24 hours. cCpE and COP-1 were added to the apical and basolateral compartments separately, while CpE, COP-2 and buffer only were added to the apical compartment only. All proteins were dialyzed in Tris buffered saline and 0.2 µm filtered before use. TEER was measured using the Millicell-ERS (Electrical Resistance System) epithelial volt-ohmmeter (Millipore Sigma) and STX2 chopstick electrode (WPI) and normalized by the surface area of the monolayer. TEER values were calculated using the following equation: (Cell resistance – Blank resistance) (ohms) x membrane surface area (cm^2^). TEER values were obtained before treatment with proteins and after overnight incubation with proteins. For the COP-1 concentration dependent assay, 200 to 9000 nM COP-1 was added to the apical compartment of Caco-2 monolayers and TEER was recorded as described above using the STX2 chopstick electrode. After measuring, media containing COP-1 was removed from the monolayers and replaced with fresh media then TEER was measured again after 24 hours.

TEER measurements from cell cultures treated as above were also measured using a modified Ussing chamber (Physiologic instruments). After treatment with test proteins, Caco-2 monolayers were removed from the transwell plate, mounted in the Ussing chamber and bathed in a ringer pH 7.4 buffer solution containing 119 mM NaCl, 21 mM NaHCO_3_, 2.4 mM K_2_HPO_4_, 0.6 mM KH_2_PO_4_, 1.2 mM MgCl_2_, 1.2 mM CaCl_2_, 10 mM glucose, at 37°C in the presence of 5% CO_2_. Basal TEER readings of each monolayer in the ringer solution was then recorded for about 2 minutes.

### Cryo-EM

Claudins purified as described above were exchanged from DDM to 2,2-didecylpropane-1,3-bis-β-D-maltopyranoside (LMNG) detergent via a PD-10 column (Bio-Rad). COP-1 was added in 1.4 moles excess to claudins then 1.4 moles excess Nb to COP-1 were mixed and incubated at 4°C for 1 hour. Complexes were concentrated, filtered, and ran on a Superdex 200 increase (5/150) gl (Cytiva) column equilibrated in 20 mM Hepes 7.4, 100 mM NaCl and 0.003% LMNG. Peak fractions were pooled and concentrated to 4 mg/mL.

3.5 μL of complex were applied to UltrAuFoil 1.2/1.3 300 mesh (Quantifoil) grids that were glow-discharged for 60 seconds at 15 mA using a Pelco easiGlow (Ted Pella Inc) instrument. Grids were blotted for 5 seconds then plunge frozen into liquid ethane cooled by liquid nitrogen using an EM GP2 (Leica) plunge freezer at 4°C and 100% relative humidity. Grids were stored in liquid nitrogen then imaged using either a 200 or 300 kV cryo-TEM.

For the receptor COP-1/Nb complex, data collection was performed on a Titan Krios G3i (ThermoFisher) equipped with a Gatan K3 direct electron detector and BioQuantum GIF at the University of Chicago Advanced Electron Microscopy Core Facility (RRID:SCR_019198). 6,774 movies were collected using EPU (ThermoFisher) in CDS mode at 105,000× magnification with a super resolution pixel size of 0.827 Å and physical pixel size of 1.65 Å, defocus range of -0.9 to - 2.1 μm with a total dose of 70 electron/Å^2^. For the non-receptor COP-1/Nb complex, data collection was performed at the Pacific Northwest Cryo-EM Center (PNCC) on a Titan Krios G3i (ThermoFisher) equipped with a Gatan K3 direct electron detector and BioContinuum HD GIF. 4,801 movies were collected using SerialEM in counting mode at 130,000× magnification with a physical pixel size of 0.649 Å and super resolution pixel size of 0.324 Å, defocus range of -0.8 to - 2.4 μm with a total dose of 50 electrons/Å^2^. For the partial receptor COP-1/Nb complex, data collection was performed at the Hauptman-Woodward Medical Research Institute (HWI) on a Glacios 2 (ThermoFisher) equipped with a Falcon 4i direct electron detector. 1,019 movies were collected using EPU (ThermoFisher) at 120,000× magnification with a physical pixel size of 0.884 Å, defocus range of -0.4 to -1.8 μm with a total dose of 50 electrons/Å^2^.

Micrograph and particle processing were performed in CryoSPARC (*71*). After patch-motion correction and patch-CTF correction of micrographs, blob-based particle picking followed by template-based picking yielded particles of complexes. These particles were then subjected to multiple rounds of 2D classifications, followed by *ab initio* 3D reconstruction and non-uniform refinement to yield the final maps (**Fig. S6**). Using the claudin/COP-1/Nb portion of PDB ID 8u4v we used Chimera to grossly fit the model into each map (*72*). No further model building or refinement of structures was conducted as the maps were not high resolution enough to visualize any protein features past secondary structure.

## Statistical analysis

Values from independent experiments are shown as mean ± standard error of mean. Statistical analysis for relative TEER measurements were performed by one-way ANOVA and Dunnett’s multiple comparison test. For statistical analysis comparing different protein treatments before and after treatment, paired student t-test analysis was used for the two-group comparison using GraphPad Prism, version 10 (GraphPad, San Diego, CA). Differences were considered significant when the P value was less than 0.05.

## Acknowledgments

We are grateful to the University of Chicago Advanced Electron Microscopy Core Facility (RRID:SCR_019198), to Craig Yoshioka, Claudia López, and Sean Mulligan at PNCC, and to Devrim Acehan and Katherine Spoth at HWI Cryo-EM Center for cryo-EM access. A portion of this research was supported by NIH grant U24GM129547 and performed at the PNCC at OHSU and accessed through EMSL (grid.436923.9), a DOE Office of Science User Facility sponsored by the Office of Biological and Environmental Research.

## Funding

Research reported in this publication was supported by the National Institute of General Medical Sciences of the National Institutes of Health under Award Numbers:

NIH NIGMS R35GM138368 (AJV)

NIH NIGMS R01GM117372 (AAK)

The content is solely the responsibility of the authors and does not necessarily represent the official views of the National Institutes of Health.

Grant support, cryo-EM training, and this material is based upon work supported by the National Science Foundation Grant OIA-2131902 (AJV)

## Author contributions

Conceptualization: AJV

Methodology: CPO, MdlA, AAK, MED, AJV

Investigation: CPO, AMM, XL, SK, JC, AJV

Resources: AAK, MED

Supervision: AJV

Funding acquisition: AAK, AJV

Writing—original draft: CPO, AJV

Writing—review & editing: CPO, AJV

## Data and materials availability

All data are available in the main text or the supplementary materials.

