## Supplement for "Biophysical Basis of Paracellular Barrier Modulation by a Pan-Claudin-Binding Molecule"

Chinemerem Ogbu *et al.*

**This PDF file includes:**

Figs. S1 to S7
Tables S1 to S2
References (1 to 11)

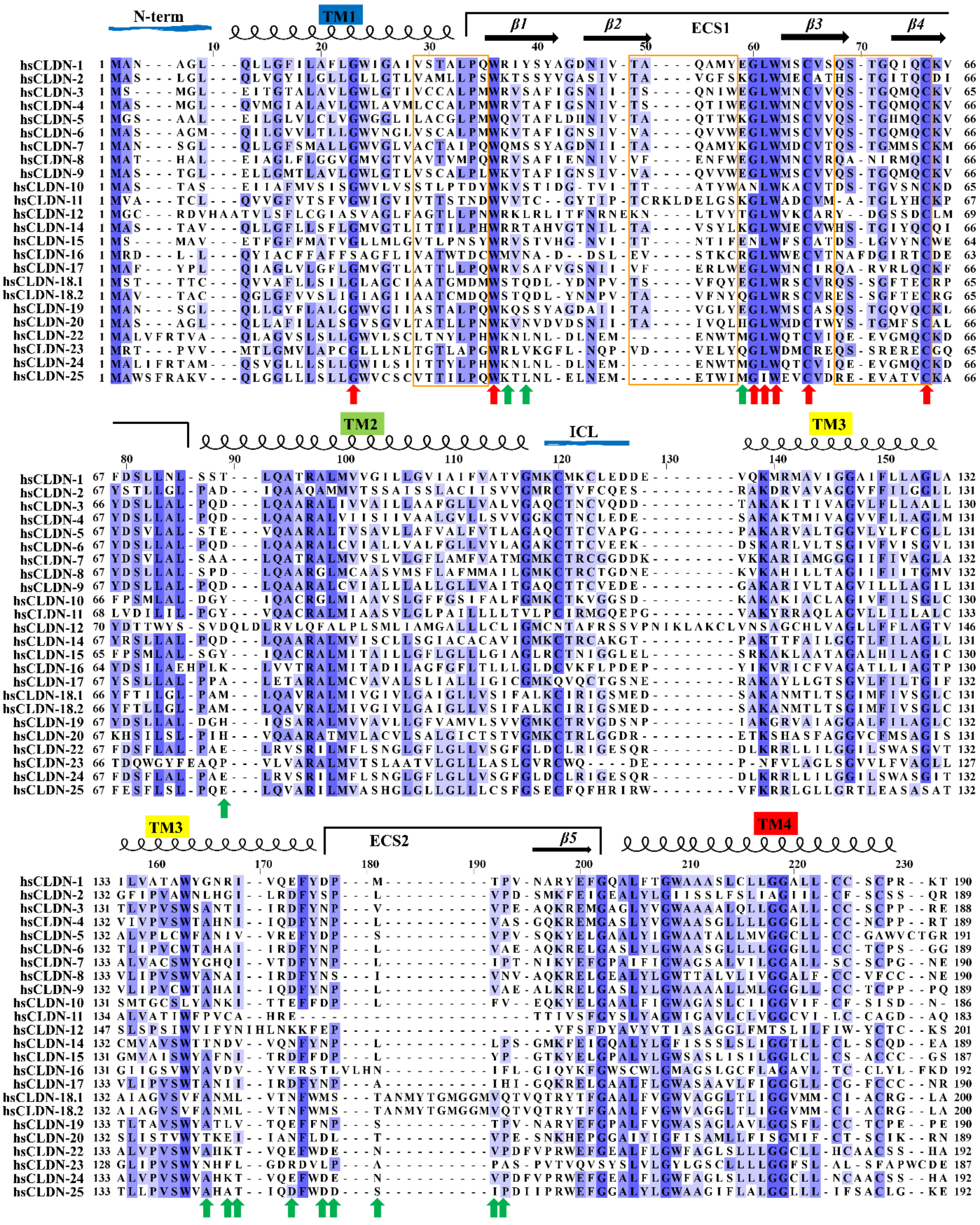


**Fig. S1. Sequence alignment of human claudins.** Sequences were aligned with T-Coffee (*1*) and visualized in Jalview (*2*). Marked above the sequences are the secondary structural elements: ECS extracellular segment; TM transmembrane domain; ICL intracellular loop. The C-terminus was aligned but is not shown above for clarity. Arrows indicate the conserved claudin family motif $W_{30}$ $GLW_{51}$ – $C_{54}$ – $C_{64}$ (red) and residues recognized by cCpE for binding to receptors (green). COP-1 binding epitopes 1, 2, and 3 are marked by boxes (orange).

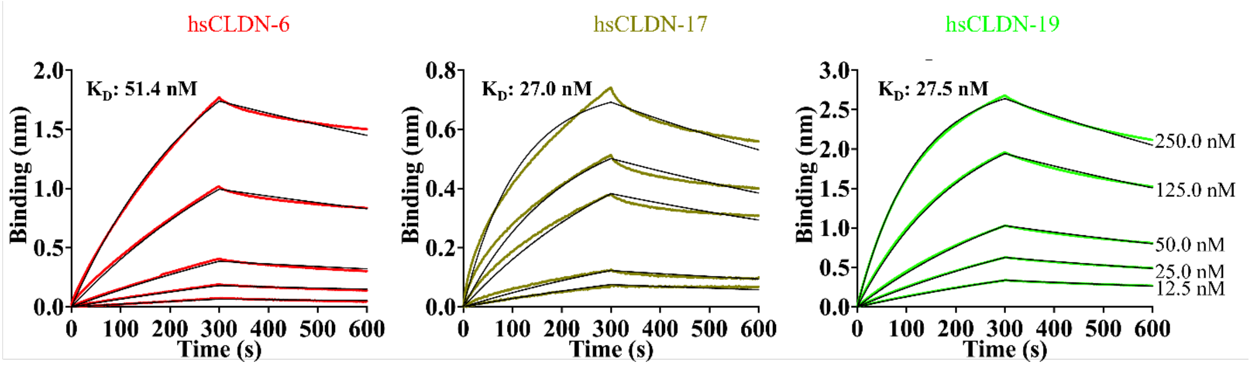


**Fig. S2. Binding of cCpE to human claudins -6, -17, and -19.** Multi-concentration point (0-250 nM) analyses of hsCLDN-6 (red), hsCLDN-17 (asparagus) and hsCLDN-19 (green) binding to immobilized cCpE using BLI. Kinetic rates appear in **Table S2**.

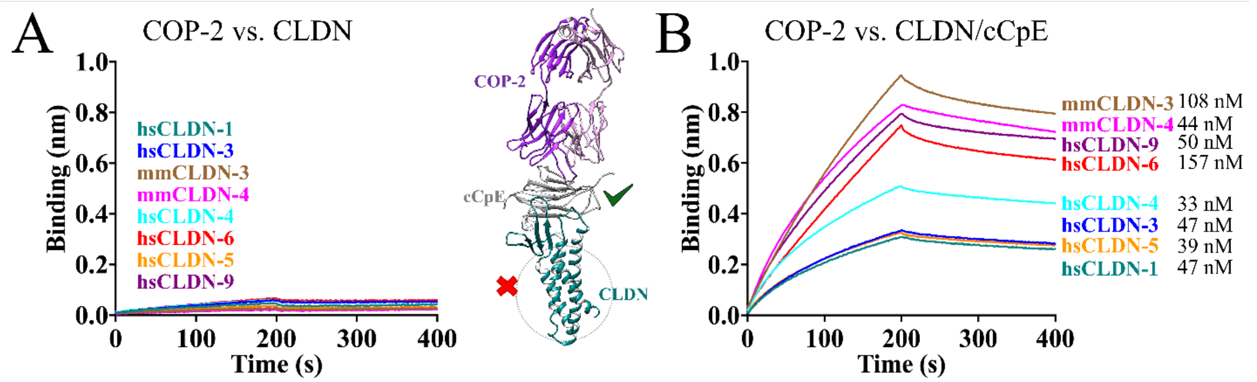


**Fig. S3. COP-2 binding to claudins or claudin/cCpE complexes.** (A) Single-concentration point (500 nM) assessment of COP-2 binding to claudins. (B) Single-concentration point (500 nM) assessment of COP-2 binding to claudin/cCpE complexes. Structural model of the expected complex between COP-2 (magenta) and claudin/cCpE (teal/grey) is shown based on PDB ID 7tdm. Note that because COP-2 binds cCpE and not claudins that non-receptors show less binding than receptors because only cCpE is being bound and not claudin/cCpE in the former.

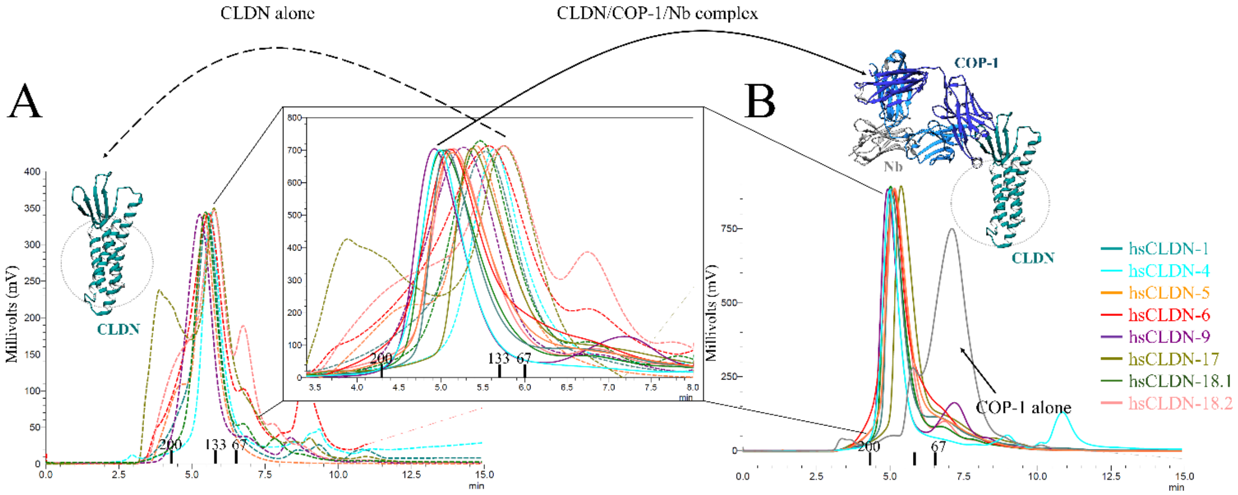


C

| Subtype | Elution time (mins) CLDN alone | Elution time (mins) +COP-1/Nb |
|---|---|---|
| hsCLDN-1 | 5.53 | 5.07 |
| hsCLDN-4 | 5.63 | 4.99 |
| hsCLDN-5 | 5.42 | 5.14 |
| hsCLDN-6 | 5.57 | 5.11 |
| hsCLDN-9 | 5.27 | 4.93 |
| hsCLDN-17 | 5.80 | 5.37 |
| hsCLDN-18.1 | 5.47 | 5.02 |
| hsCLDN-18.2 | 5.75 | 5.15 |

**Fig. S4. COP-1 binding to claudins using size exclusion chromatography.** (A) SEC chromatograms showing the elution times of claudin alone (dashed lines) and (B) claudin/COP-1/Nb complexes (solid lines). Overlays of both chromatograms are shown in the inset. Structural models of the what claudin (teal) or claudin/COP-1/Nb complex (teal/blue/grey) are in solution are shown based on PDB ID 8u4v. (C) Elution peak times of claudins versus claudin/COP-1/Nb complexes calculated from SEC.

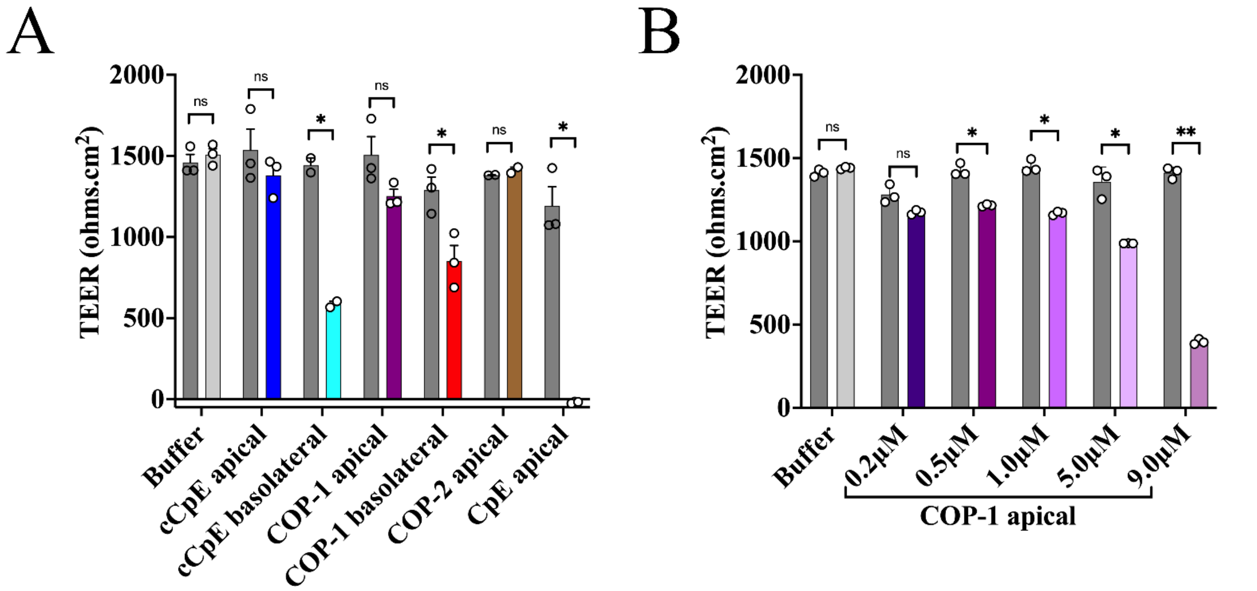


**Fig. S5. Effect of COP-1 on TEER of Caco-2 cells**. (A) TEER readings before (grey bar) and after overnight treatment (colored bar) with test proteins including cCpE (blue/cyan), COP-1 (purple/red), COP-2 (brown), and CpE (black), and buffer only control (lt. grey). TEER was measured n = 3 for buffer, COP-1 apical and basolateral, cCpE and CpE apical treatments; and n = 2 for cCpE basolateral and COP-2 treatments. Relative TEER results can be found in **Fig. 3A**. (B) TEER measurements after applying increased concentrations of COP-1 to apical compartments. n = 3 for all measurements. TEER values before treatment (grey bars) are compared to post-treatment (colored bars). T-test analysis: * represents $p < 0.05$ and ** represents $p < 0.001$. Relative TEER results can be found in **Fig. 3C**.

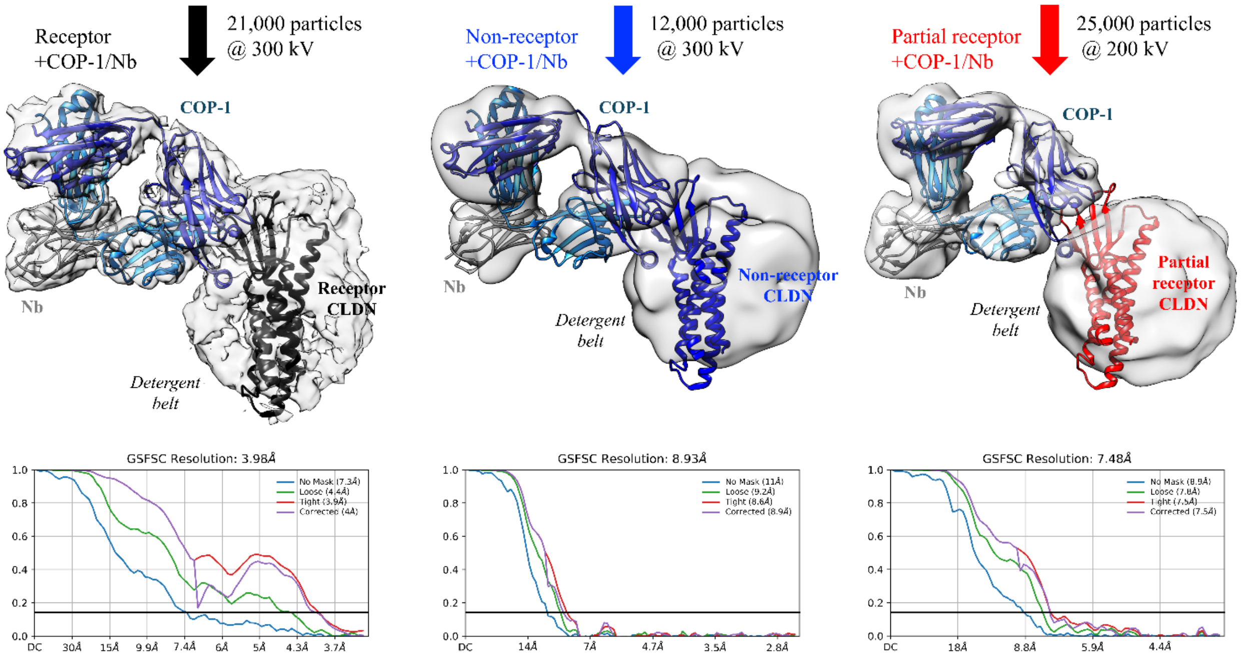


**Fig. S6. Cryo-EM map resolution of claudin/COP-1/Nb complexes.** 3D reconstructions from **Fig. 4B** are shown with corresponding gold standard Fourier shell correlation (GSFSC) curves generated by CryoSPARC at a cutoff of 0.143 **(*3*)**.

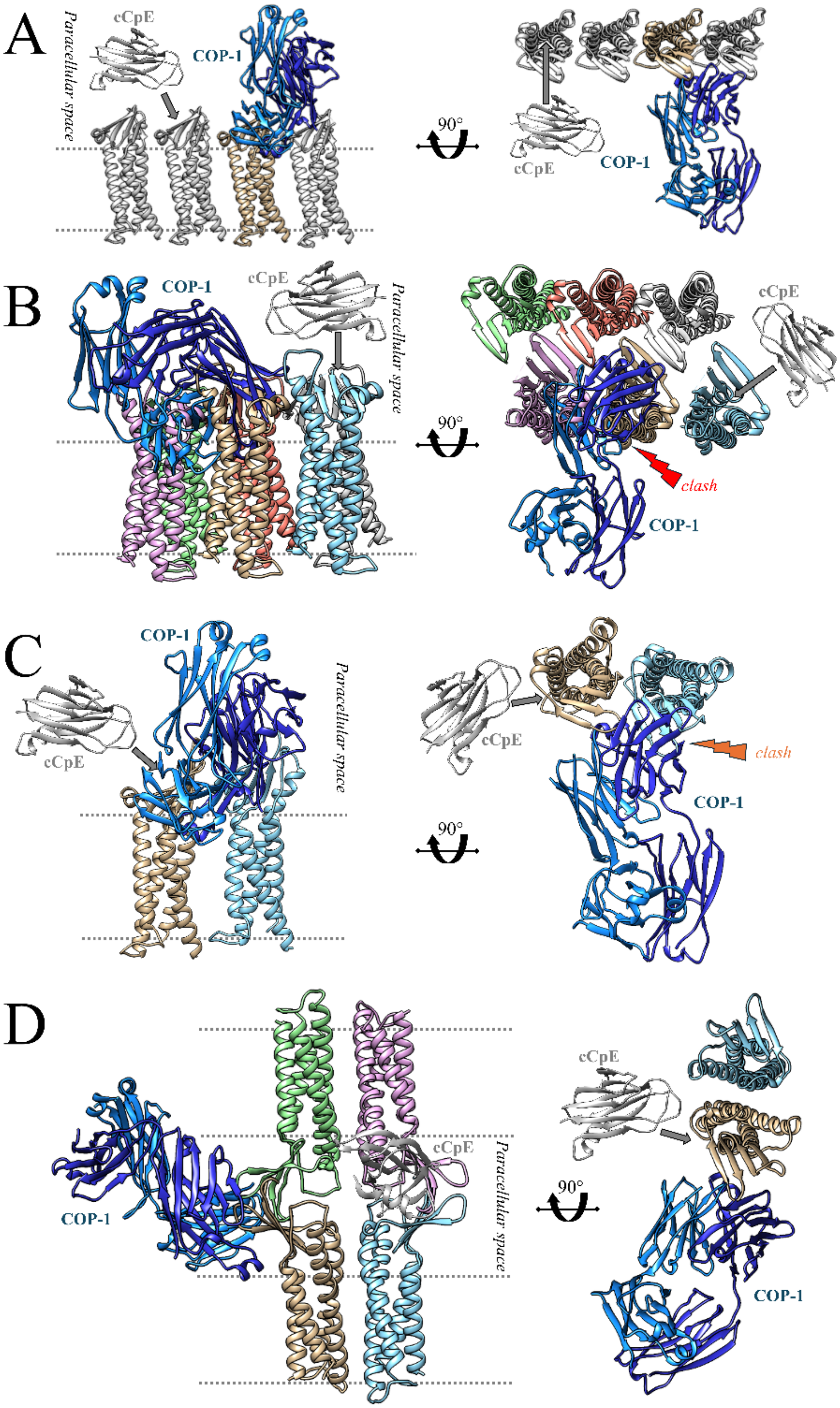


**Fig. S7. Models of polymerized claudins in tight junctions.** (A) The linear assembly of claudins as observed by Suzuki *et al.* in the crystal structure of mmCLDN-15 (***4***). (B) The face-to-face dimer (X-1) model proposed by Suzuki *et al.* (***5***). (C) The alternate cis dimer (Cis-1) proposed by Zhao *et al.* (***6***). (D) An additional alternative cis and trans interacting dimer that we propose. Each claudin protomer is colored independently. The binding sites for cCpE (grey) and COP-1 (blue) are shown with only COP-1 forming complexes for clarity, bound to a claudin (tan). For A-C only cis interactions within the same membrane are shown whereas for D, both cis and trans assemblies are modeled. Note that cCpE binds claudins in all orientations with no clashes whereas COP-1 clashes in the B and C proposed cis assemblies.

**Table S1.**

List of human claudins used for sequence alignment, the number of amino acids and their UniProt reference ID. Data was used for **Fig. 1B** and **Fig. S1**.

| Claudin | Isoform | Number of amino acids | UniProt ID |
|---|---|---|---|
| 1 | | 211 | O95832 |
| 2 | | 230 | P57739 |
| 3 | | 220 | O15551 |
| 4 | | 209 | O14493 |
| 5 | | 218 | O00501 |
| 6 | | 220 | P56747-1 |
| 7 | 1 | 211 | O95471 |
| 8 | | 225 | P56748 |
| 9 | | 217 | O95484 |
| 10 | 1 | 228 | P78369-1 |
| 11 | | 207 | O75508 |
| 12 | | 244 | P56749 |
| 14 | | 239 | O95500 |
| 15 | | 228 | P56746 |
| 16 | | 235 | Q9Y5I7 |
| 17 | | 224 | P56750 |
| 18 | 1 | 261 | P56856-1 |
| 18 | 2 | 261 | P56856-2 |
| 19 | 1 | 224 | Q8N6F1-1 |
| 20 | | 219 | P56880 |
| 22 | | 220 | Q8N7P3 |
| 23 | | 292 | Q96B33 |
| 24 | | 220 | A6NM45 |
| 25 | | 225 | C9JDP6 |

**Table S2.**
Single-point analysis of cCpE binding to claudins compared to previously published $K_{Ds}$ and kinetics of cCpE binding to hsCLDN-6, -17 and -19. Data also appears in **Fig. 1D** and **Fig. S2**

| **Single-point cCpE binding to claudins** | | | | | |
|---|---|---|---|---|---|
| **Subtype** | **$K_D$ (nM)** | **$k_{on}$ (1/Ms)** | **$k_{off}$ (1/s)** | **$t_{1/2}$ (min)** | **Published $K_D$ (nM)** |
| hsCLDN-1 | N/A | N/A | N/A | N/A | 936.2 (***7***) |
| hsCLDN-3 | 274.0 ± 1.5 | $0.3\times10^4 \pm 1.7\times10^1$ | $9.1\times10^{-4} \pm 1.7\times10^{-6}$ | 12.7 | 246.4 (***7***), 374.7 (***8***) |
| mmCLDN-3 | 12.2 ± 0.1 | $2.9\times10^4 \pm 7.4\times10^1$ | $3.5\times10^{-4} \pm 3.6\times10^{-6}$ | 33.0 | 9.4 (***7***), 9.0 (***8***), 7.9 (***9***) |
| hsCLDN-4 | 8.5 ± 0.1 | $2.7\times10^4 \pm 5.0\times10^1$ | $2.3\times10^{-4} \pm 2.6\times10^{-6}$ | 50.2 | 2.5 (***7***), 7.7 (***8***), 3.4 (***10***) |
| mmCLDN-4 | 6.4 ±0.1 | $7.0\times10^4 \pm 3.9\times10^2$ | $4.5\times10^{-4} \pm 6.3\times10^{-6}$ | 25.7 | 11.9 (***8***) |
| hsCLDN-5 | N/A | N/A | N/A | N/A | - |
| hsCLDN-6 | 52.5 ± 0.4 | $1.0\times10^4 \pm 3.6\times10^1$ | $5.1\times10^{-4} \pm 3.1\times10^{-6}$ | 22.7 | - |
| hsCLDN-9 | 4.5 ± 0.1 | $3.4\times10^4 \pm 9.7\times10^1$ | $1.5\times10^{-4} \pm 3.9\times10^{-6}$ | 77.0 | 4.9 (***8***), 3.58 (***11***) |
| mmCLDN-15 | N/A | N/A | N/A | N/A | - |
| hsCLDN-17 | 52.4 ± 0.7 | $1.1\times10^4 \pm 8.0\times10^1$ | $5.7\times10^{-4} \pm 6.8\times10^{-6}$ | 20.3 | - |
| hsCLDN-18.1 | N/A | N/A | N/A | N/A | - |
| hsCLDN-18.2 | N/A | N/A | N/A | N/A | - |
| hsCLDN-19 | 29.3 ± 0.2 | $2.1\times10^4 \pm 4.7\times10^1$ | $6.1\times10^{-4} \pm 3.0\times10^{-6}$ | 18.9 | - |

| **Binding kinetics of cCpE binding to hsCLDN-6, -17 and -19** | | | | |
|---|---|---|---|---|
| **Subtype** | **$K_D$ (nM)** | **$k_{on}$ (1/Ms)** | **$k_{off}$ (1/s)** | **$t_{1/2}$ (min)** |
| hsCLDN-6 | 51.4 ± 0.2 | $1.2\times10^4 \pm 4.3\times10^1$ | $6.1\times10^{-4} \pm 1.8\times10^{-6}$ | 18.9 |
| hsCLDN-17 | 27.0 ± 0.2 | $3.3\times10^4 \pm 1.3\times10^2$ | $8.8\times10^{-4} \pm 3.9\times10^{-6}$ | 13.1 |
| hsCLDN-19 | 27.5 ± 0.5 | $3.1\times10^4 \pm 3.6\times10^1$ | $8.4\times10^{-4} \pm 1.1\times10^{-6}$ | 13.8 |
